# Effects of tree plantations on population and roosting ecology of an endemic agamid lizard in Western Ghats Biodiversity Hotspot

**DOI:** 10.1101/2025.02.10.637441

**Authors:** Ninad Gosavi, Himanshu Lad, Jithin Vijayan, Rohit Naniwadekar

## Abstract

Low-elevation wet tropical forests support herpetofaunal diversity but are increasingly replaced by monoculture tree plantations and have poor Protected Area coverage, leaving herpetofauna vulnerable. Research on how monoculture plantations affect reptile populations remains inconclusive, and the ecology of endemic species in these changing landscapes is poorly understood. We compared densities and roosting ecology of *Monilesaurus rouxii*, an endemic agamid lizard, across low- and high-elevation forests and cashew and rubber plantations in the Western Ghats Biodiversity Hotspot using nocturnal transects. We found that low-elevation forests, despite poor Protected Area coverage, had significantly higher densities of *Monilesaurus rouxii* than high-elevation forests and cashew plantations. Roost site use did not differ significantly across the different land-use, indicating roost fidelity. However, *Monilesaurus rouxii* exhibited ontogenetic shifts in roosting substrate and height. Our findings highlight the conservation importance of unprotected low-elevation forests, which are rapidly being converted to monoculture tree plantations. We show that plantation types may differ in their impacts on reptile populations. We also show the utility of nocturnal transects for population estimation of diurnal lizards.

## INTRODUCTION

Human modification of tropical forests is a major driver of biodiversity loss worldwide, particularly impacting terrestrial vertebrates (Laurance, 2004; Phillips et al., 2017; Powers & Jetz, 2019). While agroforestry is recommended as a sustainable alternative for mitigating forest loss that is primarily driven by increasing demand for food production (FAO 2022; 2024), studies show that expanding agroforestry can impact species composition, diversity, and abundance (Boinot et al., 2022; Bohada-Murillo et al., 2020; Ferreira et al., 2020). Reptiles, despite being the most vulnerable vertebrate groups globally, have received scant attention in studies comparing different land uses (Cervantes-López & Morante-Filho, 2024; Palacios et al., 2013; Jithin et al., 2024). Given that land-use change can alter microhabitat conditions, which are critical for cold-blooded reptiles, it is crucial to determine how different forms of plantations impact their populations.

While a recent meta-analysis demonstrated that native reptile abundance was lower in monoculture plantations in the tropics (López-Bedoyaa et al. 2022), another did not find any effect of plantations on squamate abundance (Doherty et al. 2020). Most studies included in the meta-analysis were from the Americas, with poor representation from the Asian tropics, which harbour distinct reptilian lineages. Additionally, reptile species can respond differentially to plantation types as plantations might differ in the quality and quantity of microhabitats (Lad et al. 2024). For example, reptile abundance did not differ significantly between primary forests and cacao plantations in Sulawesi (Wanger et al. 2009), whereas both the endemic and wide-ranging reptile species in the Western Ghats, the white-striped viper gecko (*Hemidactylus albofasciatus*) and the saw-scaled viper (*Echis carinatus*), were less frequently seen in monoculture mango and cashew tree plantations (Jithin et al. 2023). These inconsistencies underscore the need for focused studies assessing species-specific responses to different plantation types to identify biodiversity-friendly tree plantations. This is especially critical for endemic species with smaller geographic ranges, distinct evolutionary histories, and specialised niche requirements and are more threatened than non-endemic species.

Temperature is a significant factor influencing reptilian distribution (McCain 2010), playing a critical role in determining species ranges along elevational gradients. For example, the endemic Anaimalai spiny lizard (*Salea anamallayana*) is restricted to the high-elevation evergreen forests highlighting its climatic niche fidelity (Deepak & Vasudevan, 2008). On humid forested mountains, reptilian diversity peaks at lower elevations (McCain 2010), yet Protected Areas are disproportionately concentrated in high-elevation areas (Jenkins & Joppa, 2009; Elsen et al. 2018). This mismatch suggests current conservation efforts may not safeguard key reptilian populations. Thus, understanding species responses to elevation is essential for evaluating the effectiveness of Protected Areas for conserving reptiles.

In addition to habitat selection, it is also critical to understand the microhabitat use of reptiles, especially for critical behaviours like roosting. Although sleep is essential for maintaining physiological and cognitive processes (Rattenborg et al., 2017), it remains a largely under-studied topic, particularly in wild populations (Aulsebrook et al., 2016). The selection of microhabitats for the sleeping perch is likely driven by predation avoidance, thermoregulation, and metabolic cost reduction during sleeping in reptiles (Christian et al., 1984; Libourel & Herrel, 2016). Experimental studies suggest that intraspecific interactions (Delaney & Warner, 2017) and age-sex differences (Barnett & Peo, 2023) influence sleep site selection. Thus, sleep behaviour is expected to undergo selection pressure and evolutionary adaptation, with individuals selecting more secure and thermally suitable sleep sites having a greater likelihood of survival and reproductive success (Lima & Dill 1990). Unfortunately, herpetofauna is poorly researched regarding roosting aspects (but see Libourel & Herrel, 2016; Mohanty et al., 2022). Moreover, information on age-related differences in roosting behaviour is lacking for reptiles. Understanding these differences can provide insights into age-specific predation risks and competition for roosting sites.

Given this background, we investigated the responses of Roux’s Forest Lizard (*Monilesaurus rouxii*) to different land-use categories and examined its roosting ecology. *Monilesaurus rouxii* is an agamid lizard endemic to the Western Ghats portion of the Western Ghats-Sri Lanka Biodiversity Hotspot, with only four known species in the genera (Pal et al., 2018). The Western Ghats are known for high diversity and endemism of herpetofauna, yet have faced significant forest conversion to monoculture plantations (Reddy et al. 2016). Given that protected areas are disproportionately distributed in higher elevations, it is critical to determine whether low-elevation forests and plantations are suitable for the endemic lizard species. We specifically compared the population densities of adults and juveniles *Monilesaurus rouxii* across four land-use categories⸺low- and high-elevation forests, cashew plantations and rubber plantations⸺using Distance Sampling. Additionally, we examined differences in roosting ecology between adults and juveniles. Since monoculture plantations like cashew and rubber have simplified forest structures with reduced understorey cover (Lad et al. 2024) and lower invertebrate abundances (Joshi, 2020), we hypothesized that *Monilesaurus rouxii* densities would be significantly lower in these habitats. Given that other vertebrates, like owls (Bock et al. 2013) and bats (Linton & MacDonald, 2019), exhibit age-related differences in roost site preferences, we expected adult and juvenile *Monilesaurus rouxii* lizards to differ in the use of roost sites. We find that low-elevation forests have the highest densities of lizards, and we find ontogenetic differences in roost use. We also demonstrate the use of nocturnal line-transect surveys for estimating agamid lizard densities in tropical forests.

## METHODS

### Study area

We conducted the study in the northern portion of the Western Ghats in the Sindhudurg District of Maharashtra, India (15°40’–16°0’N; 73°51’–74°9’E). Western Ghats is one of the “hottest” biodiversity hotspots because of high endemism, especially of herpetofauna, and the threats it faces (Luedtke et al., 2023). Ramachandra et al. (2017) documented a significant increase in plantation cover, within the Sahyadri region of the Western Ghats during the period from 2004 to 2013. While many new endemic species of herpetofauna continue to be described from the region, little is known about the impacts of human activities such as plantations on these endemic species (Pal et al. 2018). Most of the Protected Areas are restricted to higher elevations in the northern Western Ghats, and apart from a few patches of government-owned Reserved Forests, most forests in the region are privately owned. These forests are being converted into rubber and cashew plantations at an alarming rate in the lower elevations (Rege & Lee 2022; Munje & Kumar 2022; Jithin et al. 2023; Lad et al., 2024; Fig. 1A).

**Figure 1.**
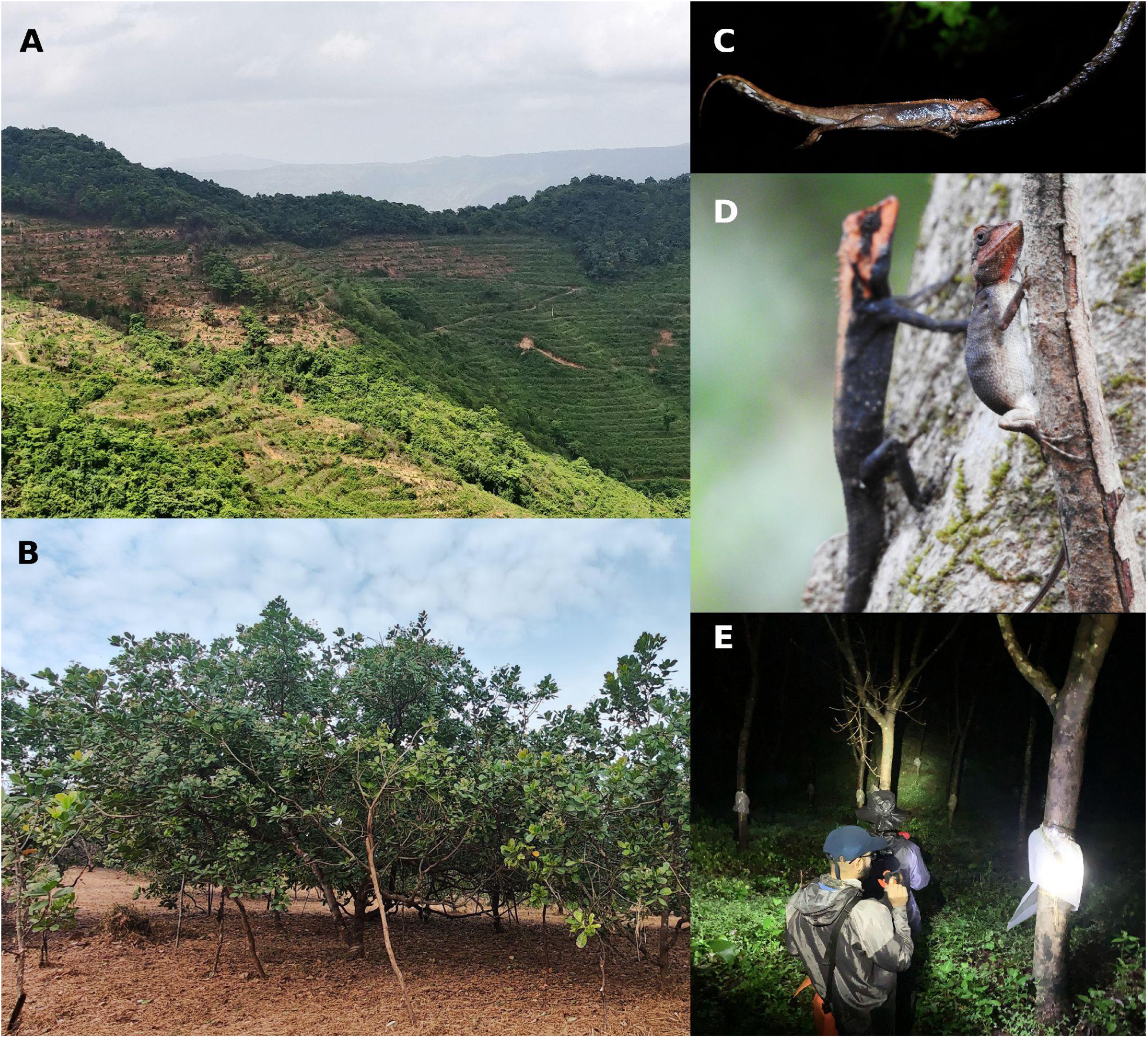
Photographs depicting (A) large-scale forest conversion to cashew plantations in the northern Western Ghats; (B) cashew monoculture plantation; (C) a roosting *Monilesaurus rouxii* in inward position, shining by the torchlight; (D) male and female *M. rouxii*; and (E) NG and HL conducting a nocturnal transect in a rubber plantation. Photograph credits: Rohit Naniwadekar and Ninad Gosavi.

### Field methods

We established 30 line transects of variable lengths across the four land-use types, including the low- and high-elevation forests (Table S1). Two observers (NG and HL) walked these transects in the night from 1830 - 2330 hr following the distance sampling survey methodology (Buckland et al. 2005; Fig 1D). We systematically searched for *Monilesaurus* lizards using a hand-held torch. Whenever a roosting *Monilesaurus* was detected, we recorded the perpendicular distance of the animal from the transect and the height of the roosting animal from the ground using a distometer (Leica Geosystems D1 laser distance metre; range: 0.2–40 m; accuracy: ± 2 mm). We classified all *Monilesaurus* larger than 50 mm (Total Length) as adults, and the rest as juveniles, following Pal et al. (2018). We also recorded the orientation of the roosting lizards as inwards, outwards, upwards, or downwards, following established methods (Mohanty *et al*. 2016, Bors et al. 2020). In addition to this, we recorded the roosting substrate, which included climber, shrub (leaf, stem, primary or secondary branch), grass (leaf), and tree (secondary branches).

### Analyses

All analyses were carried out in R 4.3.3 (R Core Team, 2021). We used multi-covariate distance sampling (MCDS) implemented by package ‘distance’ in R to estimate densities of *Monilesaurus rouxii* across different land-use types and age groups (Buckland *et al*. 2005; Marques et al. 2007; Miller et al. 2019). We used the standard key functions (half-normal and hazard rate) and right-truncated 5% of extreme distances. Since we expected variation in the detection probabilities, we modelled detection probability as a function of land-use type and age-class. We used a generalised linear model with negative binomial error structure to examine the relationship between the roost height of the lizards as a function of land-use category and age.

We used Chi-square test of independence to see if substrate use or orientation during roosting differed across land-use and age-class categories.

## RESULTS

### Density

We had 187 detections of *M. rouxii* across different land-use types (adults = 100; juveniles = 87). The hazard-rate model fitted the data well (CvM *χ*^2^ statistic = 0.17; GoF-*p* = 0.32).

Detection probability of *M. rouxii* differed across land-use types and age classes (Table S1). The overall detection probability was high (mean ± SE: 0.736 ± 0.043). *M. rouxi* density was higher in low-elevation forests than in high-elevation forests (Fig. 2). The density was lower in cashew plantations compared to low-elevation forests (as inferred by 95% CI not overlapping the means) (Fig. 2A). The density of individuals between low-elevation forest and rubber plantations, and high-elevation forest and cashew plantations did not differ (as inferred by 95% CI overlapping the means (Fig. 2A). Interestingly, the overall densities of adults and juveniles were similar (Fig. 2B).

**Figure 2.**
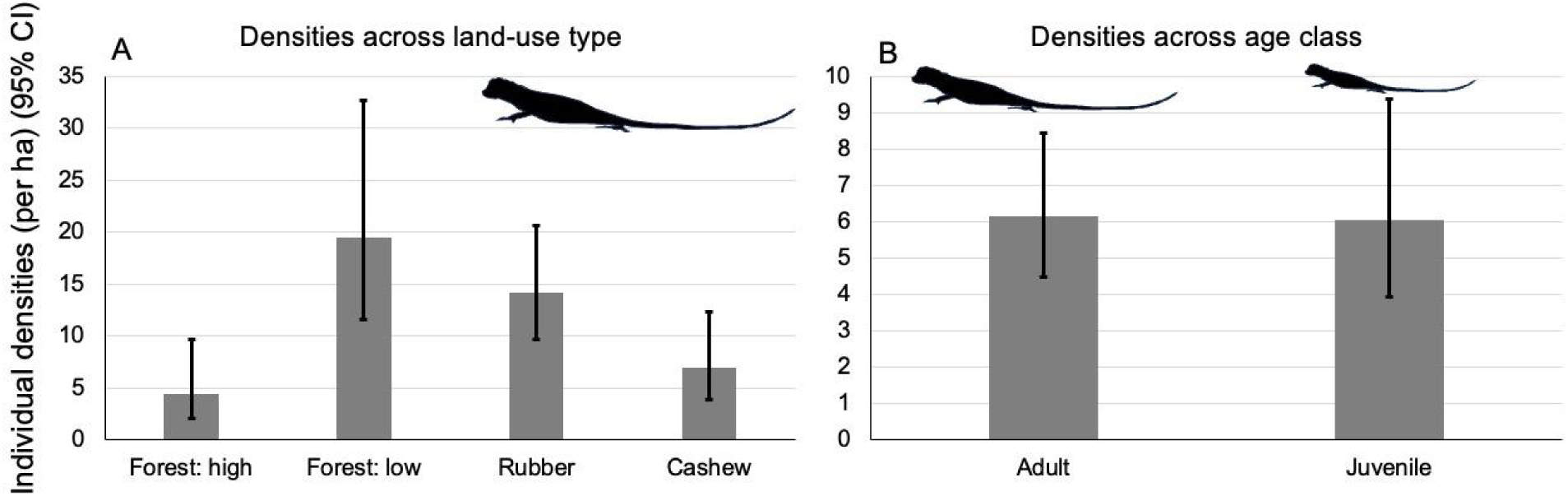
Bar plots showing the individual densities (per ha) (± 95% CI) of *Monilesaurus rouxii* across different (A) land-use types and (B) age classes. Lizards with total length ≥ 50 mm were considered adults, and < 50 mm were considered juveniles. Cashew and high-elevation forests had significantly lower densities than low-elevation forests.

### Roosting height

The roosting height of adult *M. rouxii* varied between 152.5–180.6 cm across different land-use types, while that of juveniles varied between 39–95.3 cm (Table S2). We did not find a statistically significant difference in the height of the roosts of *M. rouxii* across land-use types (Table S3). The roosting height of adult *M. rouxii* was between 1.6–4.2 times higher than juveniles (Table S2). However, juvenile *M. rouxii* roosted at lower heights than adults (Fig. 3; Table S3).

**Figure 3.**
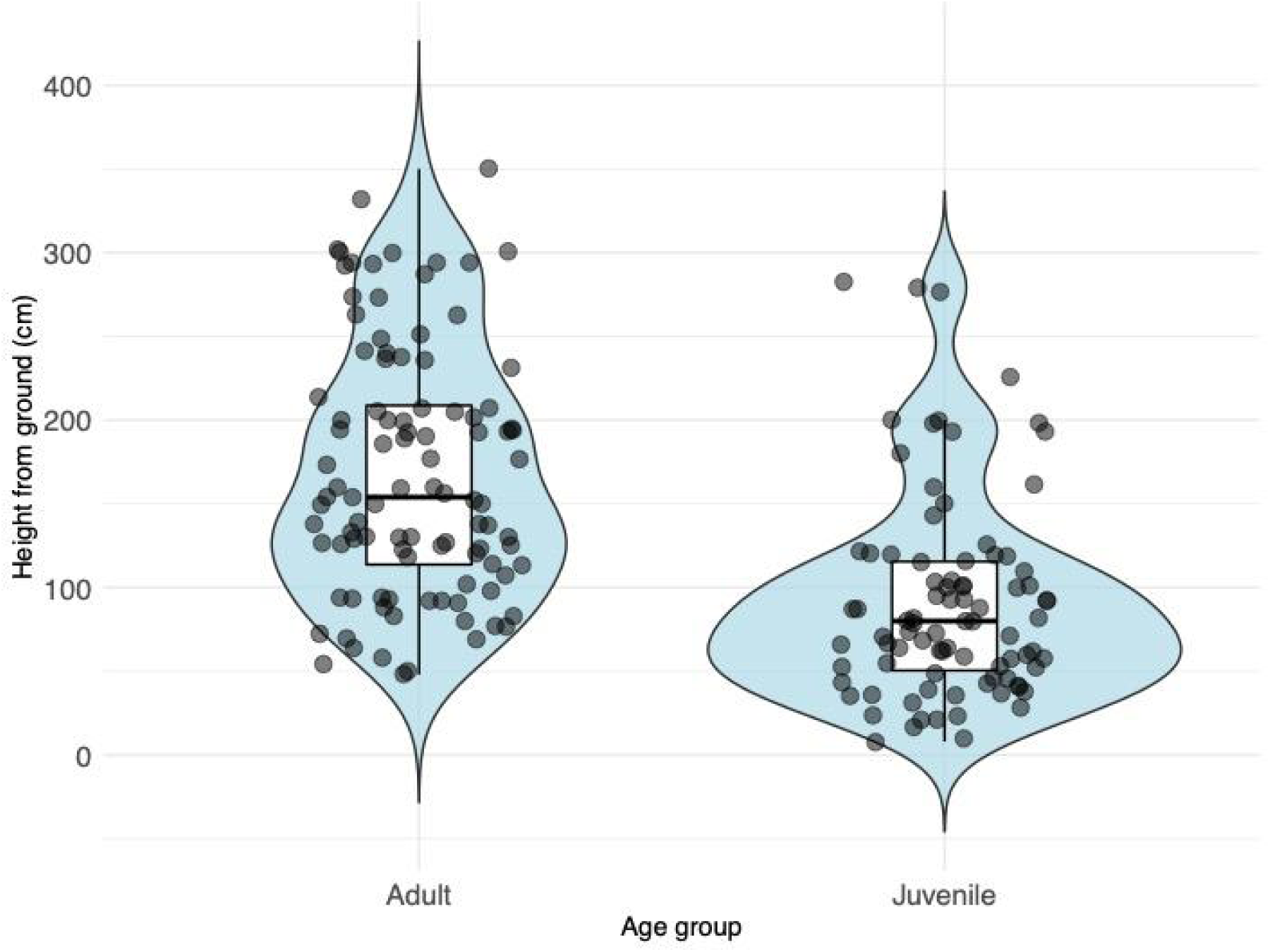
Violin plots showing the distribution of roosting heights (cm) of adult and juvenile *Monilesaurus rouxii*. The violin plots show the combined visualisation of boxplots and density traces. Points are individual data points.

### Roosting substrate and orientation

We found that the substrate used for roosting differed between adults and juveniles (*χ^2^* = 79.35, df = 6, *p* < 0.001). Adults mostly used primary branches of shrubs, followed by secondary branches of trees and shrubs to roost (Fig. 4A). However, the juveniles mostly used the main stem and leaves of shrubs to roost (Fig. 4A). We did not detect adult *M. rouxii* using leaves of shrubs or grasses for roosting during our sampling. We also found that the orientation while roosting differed across adults and juveniles (*χ^2^* = 25.35, df = 3, *p* < 0.001). Adults mostly faced inwards towards the main stem/trunk followed by outward orientation (Fig. 4B). However, the orientation of juveniles was mostly inwards or downwards (Fig. 4B). For adults, we did not find differences in substrate use (*χ^2^* = 16.11, df = 12, *p* = 0.186) or orientation (*χ^2^* = 6.74, df = 9, *p* = 0.66) across land-use types. For juveniles, we did not find differences in substrate use (*χ^2^* = 21.36, df = 18, *p* = 0.262) or orientation (*χ^2^*= 16.75, df = 9, *p* = 0.053) across land-use types.

**Figure 4.**
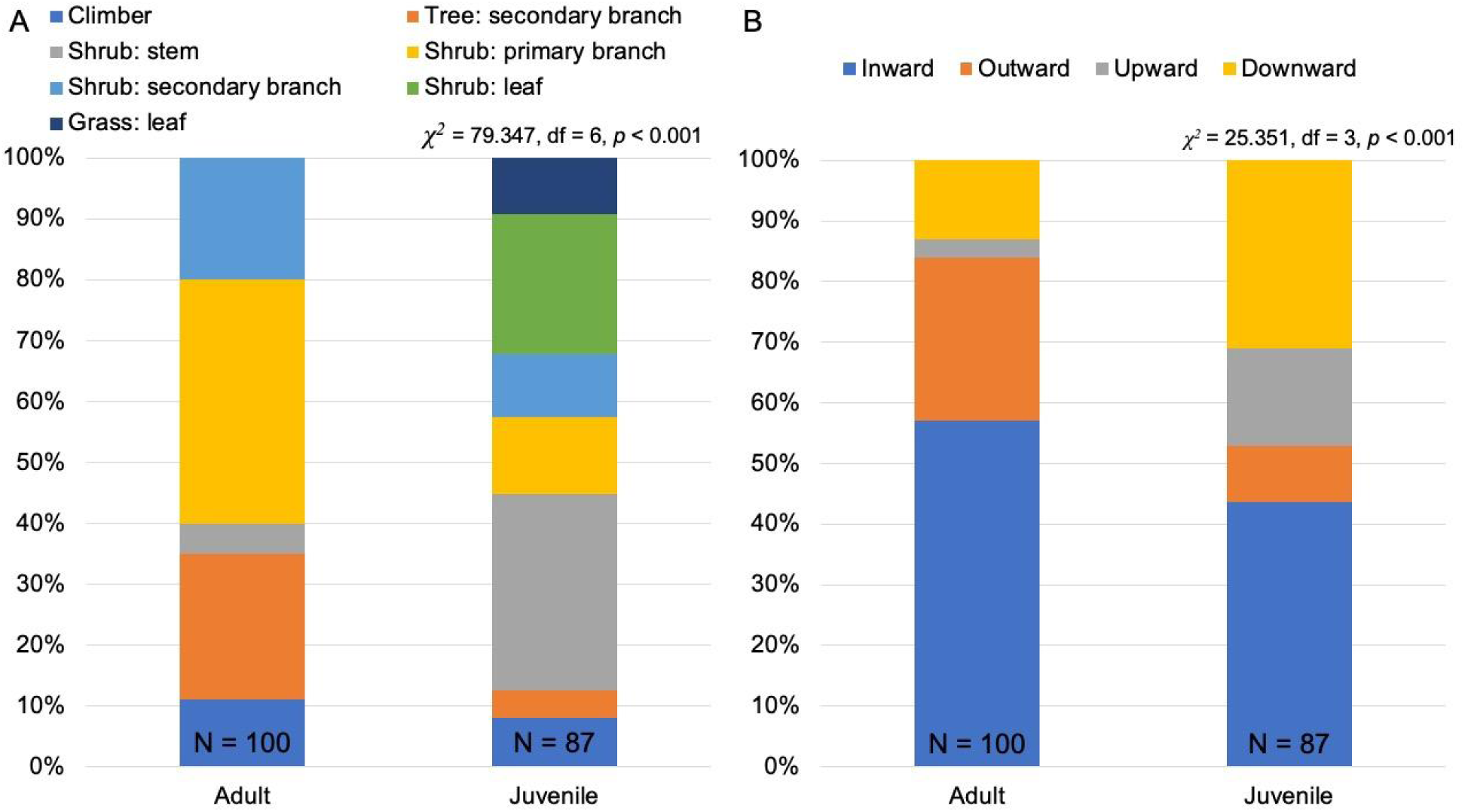
Stacked bar plots showing the percentage of roosting adult and juvenile *Monilesaurus rouxii* detections across different (A) substrates and (B) roosting orientations. The total number of *M. rouxii* adult and juvenile observations is reported at the bottom of each bar. The adult and juvenile *M. rouxii* differed in the substrate use and orientation while roosting.

## DISCUSSION

We compared the densities and roosting ecology of the endemic lizard *Monilesaurus rouxii* across multiple land-use types in the Western Ghats, a biodiversity hotspot significantly impacted by tree monoculture plantations. We found that *Monilesarus rouxii* densities were highest in low-elevation forests, while they were lower in cashew plantations and in high-elevation forests, underscoring the importance of low-elevation forests for this species. While roost site use did not differ across land-use types, juveniles and adults exhibited significant differences in roosting heights, orientation, and substrate use. We build on previous studies, by demonstrating the use of night transects for estimating densities of diurnal, arboreal lizards and comparing the roosting ecology of adults and juveniles.

### Importance of low-elevation forests for *Monilesaurus rouxii*

In humid tropical forests, reptile diversity is highest in low-elevation forests, influenced by favourable temperatures and more cloud-free days compared to high elevation areas (McCain et al. 2010; Jins et al. 2022). Our results align with this pattern, showing significantly higher densities of *Monilesaurus rouxii* in low-elevation forests. This finding is consistent with other studies from the region, which have highlighted the importance of low-elevation forests for biodiversity, including endemic bush frogs (Lad et al. 2024) and birds (Biswas et al., 2023).

Despite their ecological significance, low-elevation forests are poorly represented within Protected Areas, a pattern documented across the tropics (Jenkins & Joppa, 2009; Elsen et al. 2018). In northern Western Ghats, most Protected Areas are located at higher elevations, with no Protected Areas in the Ratnagiri and Sindhudurg Districts of Maharashtra (Kulkarni & Mehta, 2013), which harbour some of the last remaining low-elevation forests of northern Western Ghats. Meanwhile, privately-owned forests are rapidly being converted into agroforestry plantations, particularly cashew plantations, which now cover a third the land in two sub-divisions of the district of our study area (Rege & Lee, 2022). Given that *Monilesaurus rouxii* densities in cashew plantations were nearly a third of those in low-elevation forests, it is critical to preserve existing forest patches and restore degraded ones through partnerships with local communities.

### Differential responses of *Monilesaurus rouxii* to different plantation types

Previous meta-analyses have reported neutral or negative impacts of plantations on reptilian abundances in the tropics (Doherty et al. 2020; López-Bedoya et al. 2022). Our study found significantly lower densities in cashew plantations compared to low-elevation forests, but densities in rubber plantations were not significantly different from those in forests. This pattern is similar to our previous findings on *Pseudophilautus amboli*, an endemic shrub frog (Lad et al. 2024).

Agamid lizards prefer wooded areas and are negatively impacted by agricultural intensification (Balouch et al. 2022). *Monilesaurus* lizards, in particular, prefer forested areas with complex vegetation (Pal et al. 2018). Our results suggest that understorey vegetation is critical for *Monilesaurus rouxii*, as they extensively used shrubs as roosts. This understorey vegetation is also likely to be critical for them for foraging and basking during the daytime. Importantly, we found no differences in substrate use across land-use types, indicating strong fidelity to understorey vegetation. Previous habitat assessment along the same transects revealed that shrub cover was relatively higher in rubber than in cashew plantations (Lad et al. 2024). This is indicative of intensive management practices and more canopy openness in cashew plantations leading to lower shrub and higher grass cover. Reduced microhabitat availability is a known driver of reptilian population declines in tree plantations, as demonstrated in oil palm plantations in Colombia (Gallmetzer & Schulze, 2015). Beyond habitat structure, prey availability can influence lizard densities. A global meta-analysis (Wang et al. 2022) and a previous study in the region (Joshi, 2020), have documented lower invertebrate abundance in plantations compared to primary forests. As tree plantations alter both biotic and abiotic parameters, future studies should examine the interactions between habitat structure, thermal environment, and food availability and their impacts on lizard physiology and population dynamics.

### Roosting ecology of adult and juvenile *Monilesaurus rouxii*

Differences in roost site use may be driven by multiple factors, including predation risk, thermoregulation and physical constraints. In Barrington Land Iguanas (*Conolophus pallidus*), juveniles chose roosting sites offering better protection from predators, regardless of thermal conditions (Christian et al. 1984). Similarly, we observed juvenile *Monilesaurus rouxii* roosting on shrub or grass leaf blades more frequently than adults, who never used these substrates.

However, juveniles were also more often found roosting on the main stem, a site that likely presents a higher predation risk due to easier accessibility for arboreal predators. The ability of juveniles to balance more easily on leaves compared to adults (Montgomery et al. 2011) may also influence substrate choice. This contrasting microhabitat use underscores the importance of vertical structural complexity in providing suitable perching options for different age groups.

*M. rouxii* adults were observed roosting at higher perchs compared to juveniles, a pattern consistent with previous studies (Razafimahatratra et al., 2008). Higher sleeping perches may reduce predation risk and have thermal advantages as higher perches will expose lizards to radiation earlier during the day as suggested in earlier study (Montgomery et al. 2011). Thus, ontogenetic differences in substrate and perch height could be a response to predation pressure, thermal environments and intraspecific interactions (Stamps, 1983; Smith & Ballinger, 2001; Delaney & Warner, 2017).

Orientation also differed between age groups. The inward orientation observed in the most adult *Monilesaurus rouxii* in our study aligns with previous research indicating enhanced visual and tactile detection of predators (Clark & Gillingham, 1990; Carbera-Guzmán & Reynoso, 2010; Mohanty et al. 2016). On the other hand, while juveniles were found on leaves with an inward orientation, they were also predominantly found on shrub stems with a downward orientation, which is intriguing as arboreal predators are likely to approach the juveniles from above.

The absence of any difference in perch height, substrate and orientation between land-use classes is contrasting to a previous study (Miller, 2017). This suggests stronger fidelity to substrate use and with lower availability of such substrates in plantations (Lad et al., 2024), lizards can be expected to be impacted by it.

### Use of night transects for estimating densities of agamid lizards

Agamid lizards are diurnal, and previous studies have estimated their densities using daytime transects (Venugopal, 2010; Deepak & Vasudevan, 2008). However, our primary interest was on roosting behaviour leading us to conduct night surveys. During reconnaissance surveys, we observed that these cryptic lizards were easily detected at night as their bodies reflected torchlight more than the surrounding environment. Moreover, the animals remained motionless after being spotted as has been documented in other studies where flight initiation distances of *Monilesaurus* lizards are much lower in nights than in daytime (Bors et al., 2020). This ensured that key distance sampling assumptions—accurate detection of animals in their original positions and measurement of perpendicular distances from the trail—were not violated. Additionally, detecting juveniles was easier at night than during the day, a hypothesis that warrants further investigation. Our estimated detection probability (0.74) of *Monilesaurus rouxii* was higher than previous estimates for *Monilesaurus rouxii* from vanilla plantations (0.4) and for *Salea anamallayana* (0.5) from forested habitats in southern Western Ghats, highlighting the efficacy of nocturnal transects for agamid density estimation. Future studies should compare detection probabilities between diurnal and nocturnal transects to assess their relative effectiveness.

### Conclusions

The study underscores the importance of low-elevation forests in a biodiversity hotspot for an endemic arboreal lizard. Our findings highlight the differential responses of *Monilesaurus rouxii* to different plantation types, likely driven by microhabitat availability. While we observed ontogenetic shifts in roost site use, we fail to find differences in roost site use across habitats indicating roost site fidelity. Future research should explore how microhabitat structure, thermal environment, and prey availability, all interact to influence lizard behaviour, physiology and populations.

## OPEN RESEARCH STATEMENT

All codes and statistical packages used in this study are cited within the manuscript will be made publicly available on acceptance of the manuscript.

## ACKNOWLEDGEMENTS

This study was funded by the On the Edge Conservation Grant (UK). We thank Maharashtra Forest Department, especially Sunil Limaye (CWLW), for giving us the necessary permits (Letter No. Desk-22(8)/WL/Research/CR-53(20-21) /3361/22-22). We thank Vishal Sadekar, Siddharth Biniwale, Praveen Desai, Parag Rangnekar, Gajanan Shetye, Vijay Karthick, Sudhakar Desai, Rohan Thakur, Hemant Ogale, and Kaka Bhise for their immense support during fieldwork. We thank Anand Osuri and Kulbhushansingh Suryawanshi for their valuable feedback and support.

## AUTHORSHIP STATEMENT

RN conceived the ideas and designed the methodology with inputs from HL and NG; HL and NG collected the data; RN and NG analysed the data; RN, NG, and VJ led the writing of the manuscript with inputs from HL. All authors contributed critically to the drafts and gave final approval for publication.

## SUPPLEMENTARY MATERIAL

**Table S1.** Summary of sampling effort across the different land-use types.

| Category | Number of transects | Transect length (range)<br>(km) | Total effort<br>(km) |
| --- | --- | --- | --- |
| Cashew | 8 | 0.28-0.30 | 9.22 |
| Rubber | 8 | 0.27-0.30 | 9.39 |
| Forest (low) | 8 | 0.30 | 7.20 |
| Forest (high) | 6 | 0.23-0.30 | 6.48 |
| **Total** | **30** | **0.23-0.3** | **32.29** |

**Table S2.**
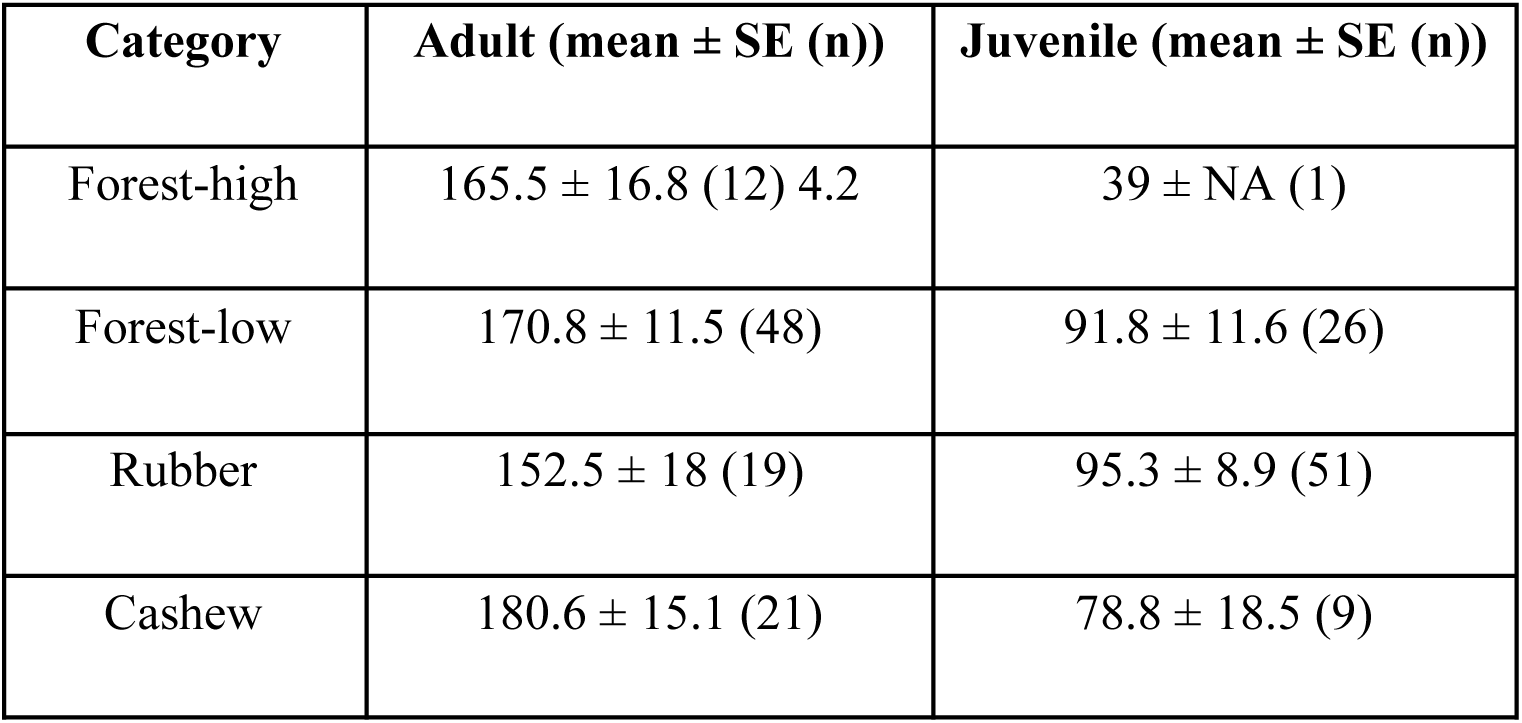
Summary of roosting heights of adult and juvenile *Monilesaurus rouxii* across different land-use types. Mean, standard error (SE) and number of observations (n) are reported. We detected only one juvenile in the high-elevation forests, therefore, we did not estimate SE for the same.

**Table S3.** Coefficient table summarising the results of generalised linear model with negative binomial error structure. The model examined the relationship between the roost height of the *Monilesaurus rouxii* lizards as a function of land-use category and age.

| **Coefficient** | **Estimate** | **SE** | ***z*** | ***p*** |
| --- | --- | --- | --- | --- |
| Intercept: Cashew & Adult | 5.137 | 0.103 | 50.031 | <0.001 |
| Land-use: Forest-high | -0.072 | 0.182 | -0.397 | 0.692 |
| Land-use: Forest-low | 0.001 | 0.118 | 0.008 | 0.993 |
| Land-use: Rubber | -0.003 | 0.125 | -0.027 | 0.978 |
| Age: Juvenile | -0.615 | 0.089 | -6.949 | <0.001 |

